# Disease stage-sensitive MFSD2A-caveolae axis insigates hippocampal blood–brain barrier disruption in Alzheimer’s disease

**DOI:** 10.64898/2026.09.30.755839

**Authors:** Yiyi Zhu, Danjie Yang, Jiayi Chen, Qi Wang, Weijie Xiong, Jingyu Wang, Hui Chen, Alexei Verkhratsky, Chenju Yi

**Author notes:** Correspondence to: Chenju Yi: No. 628 Zhenyuan Road, Guangming District, Shenzhen 518107, China;.

## Abstract

Blood-brain barrier (BBB) malfunction contributes to pathophysiology of Alzheimer’s disease (AD), yet the mechanisms that confer hippocampal endotheliocytes vulnerability remain unclear. Here we identify loss of endothelial major facilitator superfamily domain-containing protein 2A (MFSD2A), a suppressor of caveolae-mediated transcytosis, as an early and hippocampus-specific feature of AD-associated vascular impairment. In post-mortem human AD brains, reduction in hippocampal MFSD2A was accompanied with increased levels of caveolin-1 and perivascular albumin extravasation. Analysis of human single-nucleus transcriptomic data further demonstrated that vascular expression of MFSD2A declined during transition from mild cognitive impairment to established AD. In APP/PS1 mice, MFSD2A loss occurred early in hippocampal microvessels and coincided with disproportionately high hippocampal BBB leakage and reductions in GLUT1 and claudin-5, despite broader amyloid deposition throughout other brain regions. Restoration of endothelial MFSD2A using an AAV-BR1-based vascular vector during early stage of amyloid pathology, reduced albumin extravasation and preserved GLUT1 and claudin-5. Early intervention also reduced amyloid plaque burden and plaque-associated neuritic dystrophy, whereas this treatment during established amyloid pathology, although modifying selected endothelial markers, failed to reverse BBB leakage, amyloid pathology, or cognitive impairment. Sustained MFSD2A restoration from the early disease stage provided the greatest protection, preserving hippocampal BBB integrity, reducing amyloid and neuritic pathology, improving long-term potentiation and ameliorating spatial learning and memory deficits. These findings define endothelial MFSD2A loss as a disease stage-sensitive mechanism of hippocampal BBB vulnerability and highlight the MFSD2A-caveolae axis as a therapeutic target for early-stage AD.

## Introduction

Alzheimer’s disease (AD) is a progressive neurodegenerative disorder characterised by memory impairment and cognitive decline. Major neuropathological hallmarks of AD include extracellular amyloid-β (Aβ) plaques and intracellular neurofibrillary tangles ^1^. Biomarker studies suggest that Aβ pathology emerges decades before clinical manifestations, with its trajectories extending back to midlife, making it one of the earliest events in the AD continuum ^2^. Although accumulating evidence indicates that Aβ contributes to blood-brain barrier (BBB) disruption^3,4^, the underlying molecular mechanisms remain unclear, including brain region specificity of the BBB response to Aβ.

This question is closely relevant to an apparent paradox found in human brains: while the hippocampus is highly vulnerable during the early stage of AD, Aβ deposition generally emerges first in the neocortex and later extends to medial temporal regions ^5,6^. Similarly, brain-wide mapping in APP/PS1 mice showed that plaques appeared earlier and reached a greater burden in the isocortex than in the hippocampus when the disease starts ^7^. Despite the predominant cortical amyloid burden at the early disease stage, initial functional impairment mainly involves hippocampus-dependent learning and memory, whereas cortex-dependent functions remain relatively preserved. This dissociation between the anatomical distribution of amyloid and the functional deficits suggests that Aβ burden alone cannot fully explain the selective vulnerability of the hippocampus.

One potential explanation for this regional pathology-function dissociation is regional BBB heterogeneity. In individuals with early cognitive dysfunction, Dynamic Contrast-Enhanced (DCE)-MRI showed increased BBB permeability in the hippocampus, but not in the cortex ^8^. This region-specific impairment may reflect intrinsic BBB heterogeneity, as hippocampal endothelial cells exhibit distinct transport-related programmes despite conserved expression of major tight-junction genes across all brain regions ^9^. BBB integrity depends on both tight junctions and the suppression of endothelial transcytosis ^10–12^. Tight-junction disruption is the only well documented cause of BBB malfunction in AD, while the contribution of endothelial transcytosis to BBB leakage remains unknown. In several other diseases associated with BBB injury, including stroke increased caveolae-mediated endothelial transcytosis precedes junctional disruption, suggesting that it may the earliest event in compromising BBB integrity ^13^. Whether region-specific transcytosis also occurs during early AD remains unknown.

The key suppressor of caveolae-mediated transcytosis, Major Facilitator Superfamily Domain-Containing Protein 2A (MFSD2A), is enriched in the capillary endothelium of BBB-forming vessels ^10,11,14,15^. In MFSD2A knockout mice, BBB leakage persists from embryonic stage into adulthood despite normal vascular architecture and absence of overt tight-junction defects ^10^. Similarly, transport-deficient MFSD2A^D96A^ mice exhibit increased caveolae formation, endothelial transcytosis and BBB permeability, demonstrating that MFSD2A is indispensable for BBB integrity by suppressing caveolae-mediated transcytosis ^11^. In addition, MFSD2A transports docosahexaenoic acid (DHA) across the BBB in the form of lysophosphatidylcholine-DHA; loss of this function leads to brain DHA deficiency, microcephaly, neuronal loss and cognitive impairment ^14^. Despite its established roles in brain lipid transport and BBB maintenance, how MFSD2A is regulated in AD and other neurodegenerative diseases, and whether its loss contributes to disease progression, remain poorly understood. In particular, it is unknown whether vascular MFSD2A exhibits region- and stage-dependent alterations during AD progression, and whether restoring endothelial MFSD2A can protect BBB dysfunction and ameliorate neurological deficits in AD.

Here, by combining analyses of post-mortem human brain tissue, human single-nucleus transcriptomic data and age-defined APP/PS1 mice, we identified endothelial MFSD2A reduction as an early-stage and hippocampus-selective feature of AD-associated BBB dysfunction. Using three AAV-BR1-mediated MFSD2A overexpression paradigms, we found that early-stage intervention attenuated BBB malfunction and amyloid-associated pathology, whereas late-stage intervention showed only limited BBB benefits. Early and sustained MFSD2A overexpression provided the most durable protection, preserving BBB integrity while reducing neuronal pathology, enhancing hippocampal synaptic plasticity, and improving spatial learning and memory. These findings identify endothelial MFSD2A as a stage-sensitive therapeutic target in AD.

## Methods and materials

### Human brain samples

Human post-mortem brain specimens were obtained from the National Human Brain Bank for Development and Function, Chinese Academy of Medical Sciences & Peking Union Medical College under approval by the Human Research and Ethics Committee of Sun Yat-sen University (approval no. KY-2025-546-01). AD neuropathologic change was assessed by the Brain Bank in all brain specimens using the ABC scoring system and classified as Not, Low, Intermediate, or High (Supplementary Table 1).

### Mice

APP/PS1 mice expressing chimeric mouse/human APP carrying the Swedish mutation (KM670/671NL) and human PSEN1 lacking exon 9 (PSEN1ΔE9) under the control of the mouse prion protein promoter were established on a C57BL/6J background ^16^ . Mice were maintained in a PC2 animal facility with a 12 h light/dark cycle at 22 ± 2 °C with ad libitum access to food and water. All animal procedures were conducted in accordance with the NIH Guide for the Care and Use of Laboratory Animals and approved by the Animal Care and Use Committee at Sun Yat-Sen University (approval no. SYSU-IACUC-2026-002464).

### Viral vectors and virus injection

AAV-BR1-MFSD2A and the corresponding control vector (AAV-BR1-CTL) were constructed by OBIO Technology (Shanghai, China). Mouse *Mfsd2a* was cloned into a pAAV vector containing the TIE1p-mScarlet-P2A-Mfsd2a-WPRE expression cassette, in which the endothelial *Tie1* promoter drives the co-expression of an RFP reporter and MFSD2A. APP/PS1 mice were intravenously injected via the tail vein with AAV-BR1-MFSD2A (APP/PS1^MFSD2A^) or AAV-BR1-CTL (APP/PS1^CTL^) at a dose of 3 × 10¹¹ vector genomes per mouse. For early intervention, mice received a single injection at 3.5 months and were analysed at 5 months. For late intervention, mice received a single injection at 9 months and were analysed at 10 months. For sustained intervention, mice received an initial injection at 3.5 months and a second injection of the corresponding vector at 6.5 months, followed by analysis at 10 months. Mice receiving the sustained MFSD2A-overexpression or control-vector regimen were designated APP/PS1^S-MFSD2A^ and APP/PS1^S-CTL^, respectively.

### Immunofluorescence staining

Paraffin-embedded human brain sections were deparaffinised, rehydrated, and underwent antigen retrieval in citrate buffer. Sections were blocked with 10% BSA and incubated with primary antibodies diluted in 5% BSA/TBST at 4 °C for 48 h. After washing, sections were incubated with the appropriate fluorophore-conjugated secondary antibodies for 2 h at room temperature. Where indicated, Aβ plaques were labelled with Methoxy-X04 (34 μg/mL; Tocris Bioscience, #4920). Primary antibodies and labelling reagents included rabbit anti-MFSD2A (E8U2O, 1:500, Cell Signaling Technology, #80302), rabbit anti-caveolin-1 (D46G3, 1:1,500, Cell Signaling Technology, #3267S), rabbit anti-albumin (EPR20195, 1:500, Abcam, ab207327). Sections were prepared and co-stained with fluorescein-conjugated Lycopersicon esculentum lectin (#FL-1171, 1:200, Vector Laboratories) to visualise brain microvessels.

Mice were deeply anesthetised with tribromoethanol (Avertin; 250 mg/kg, intraperitoneally) and transcardially perfused with phosphate-buffered saline (PBS), followed by 4% paraformaldehyde (PFA). Brains were collected, post-fixed in 4% PFA at 4 °C overnight, and cryoprotected in 30% sucrose solution. Tissues were subsequently frozen and sectioned coronally at a thickness of 20 μm using a cryostat. Sections were rinsed in PBS and permeabilised and blocked for 1 h at room temperature in PBS containing 0.5% Triton X-100 and 2% bovine serum albumin (BSA). After blocking, sections were incubated with primary antibodies diluted in the same buffer overnight at 4 °C. Following several PBS washes, sections were exposed to the appropriate fluorescent secondary antibodies for 2 h at room temperature in the dark. Primary antibodies and labelling reagents included rabbit anti-MFSD2A (E8U2O, 1:500, Cell Signaling Technology, #80302), rabbit anti-caveolin-1 (D46G3, 1:1,500, Cell Signaling Technology, #3267S), rabbit anti-albumin (EPR20195, 1:500, Abcam, ab207327), rabbit anti-GLUT1 (1:400, Proteintech, #21829-1-AP), rabbit anti-claudin-5 (1:100, Invitrogen, #34-1600), rat anti-LAMP1 (1:200, BioLegend, #121602), rabbit anti-RTN3 (1:100, Millipore, #ABN1723). Aβ plaques were labelled with Methoxy-X04 (34 μg/mL; Tocris Bioscience, #4920). Fluorescein-conjugated Lycopersicon esculentum lectin (1:200, Vector Laboratories, #FL-1171 or #DL-1177) was used to label brain microvessels.

### Hippocampal long-term potentiation (LTP) recording

Mice were deeply anesthetised with isoflurane and transcardially perfused with ice-cold NMDG-based artificial cerebrospinal fluid (NMDG-ACSF) continuously oxygenated with 95% O and 5% CO . Brains were rapidly removed and sectioned into 400-μm thick slices using a vibratome in ice-cold NMDG-ACSF. Slices were transferred to standard ACSF, allowed to recover at 32 °C for 1 h, and subsequently maintained at room temperature for 30 min before recording. Slices were placed in a recording chamber and continuously perfused with oxygenated ACSF at 2.0-2.5 mL/min. A stimulating electrode was positioned in the Schaffer collateral pathway, and field excitatory postsynaptic potentials (fEPSPs) were recorded from the stratum radiatum of Cornu Ammonis (CA)1. Signals were amplified using a MultiClamp 700A amplifier and digitised with a Digidata 1550B system. An input–output curve was generated, and the stimulus intensity was adjusted to evoke approximately 50% of the maximal fEPSP slope. After a stable 20 min baseline, LTP was induced by delivering a train of 100 Hz (High frequency stimulation, HFS), and fEPSPs were monitored for up to 70 min after stimulation. The fEPSP slopes were normalised to the mean baseline value. Data were acquired using Clampex and analysed using Clampfit 10.7.

### Open field test

Spontaneous locomotor activity and anxiety-related behaviour were assessed using the open field test. Mice were allowed to acclimate to the animal facility for 1 week before behavioural testing. Each mouse was placed individually in a square arena and allowed to explore freely for 10 min. Locomotor activity was recorded using an automated video-tracking system. The total distance travelled, average velocity, and time spent in the central area were analysed. The arena was cleaned with 70% ethanol between trials to eliminate olfactory cues. All experiments were conducted during the light phase under consistent environmental conditions, and data were analysed by investigators blinded to the experimental groups.

### Morris Water Maze

Spatial learning and memory were evaluated using the Morris water maze. Mice were tested in a circular pool filled with opaque water maintained at 22-24 °C. A transparent escape platform was placed in a fixed location within one quadrant and submerged approximately 1 cm below the water surface. Visual cues positioned around the testing room remained unchanged throughout the experiment. During the acquisition phase, each mouse received four trials per day for five consecutive days. For each trial, the animal was released from one of several predetermined starting positions and allowed 60 s to locate the hidden platform. Animals that did not reach the platform within the allotted time were guided to it and permitted to remain there for 15 s. Escape latency, swimming distance, and swimming velocity were recorded using an automated video-tracking system. A probe test was performed 24 h after the final acquisition session. The platform was removed, and each mouse was allowed to swim for 60 s. Spatial memory performance was assessed by measuring the time spent in the target quadrant, the number of crossings over the former platform location, and the latency to first reach the platform area.

### Statistical analysis

Statistical analyses were performed using GraphPad Prism V.9. Data were assumed to follow a normal distribution. Differences between two groups were analysed using Student’s t-tests and those among more than two groups were analysed using one-way ANOVA with Tukey post hoc tests. A p value < 0.05 was considered statistically significant. Results are represented as mean ± standard error of the mean (SEM). Sample sizes were selected based on previous studies employing similar methodologies. Investigators were blinded to group allocation during data analysis.

## Results

### MFSD2A reduction and increased transcytosis in post-mortem early AD brains

We examined expression of MFSD2A in post-mortem AD brain tissues and age-matched healthy controls (Supplementary Table 1). Consistent with the diagnosis, abundant Aβ plaques were observed in the AD brains (Figure 1C, G). Before quantitative analyses, we validated MFSD2A immunostaining specificity by testing whether the observed signal reproduced two well-established features of MFSD2A cerebrovascular expression. First, MFSD2A showed the expected vessel-segment specificity^15^, with strong enrichment in capillaries in both human and mouse brains, but little to no signal in larger lectin-positive vessels (Supplementary Figure 1A). Second, MFSD2A staining was confined to BBB-forming endothelium^10,14,17^. Robust vascular staining was detected in regions with an intact BBB, whereas staining was absent from BBB-deficient vascular domains at the median eminence and arcuate nucleus interface (Supplementary Figure 1B). Thus, the staining pattern recapitulated the known capillary enrichment and BBB-associated distribution of MFSD2A, supporting the subsequent quantitative analyses.

**Figure 1.**
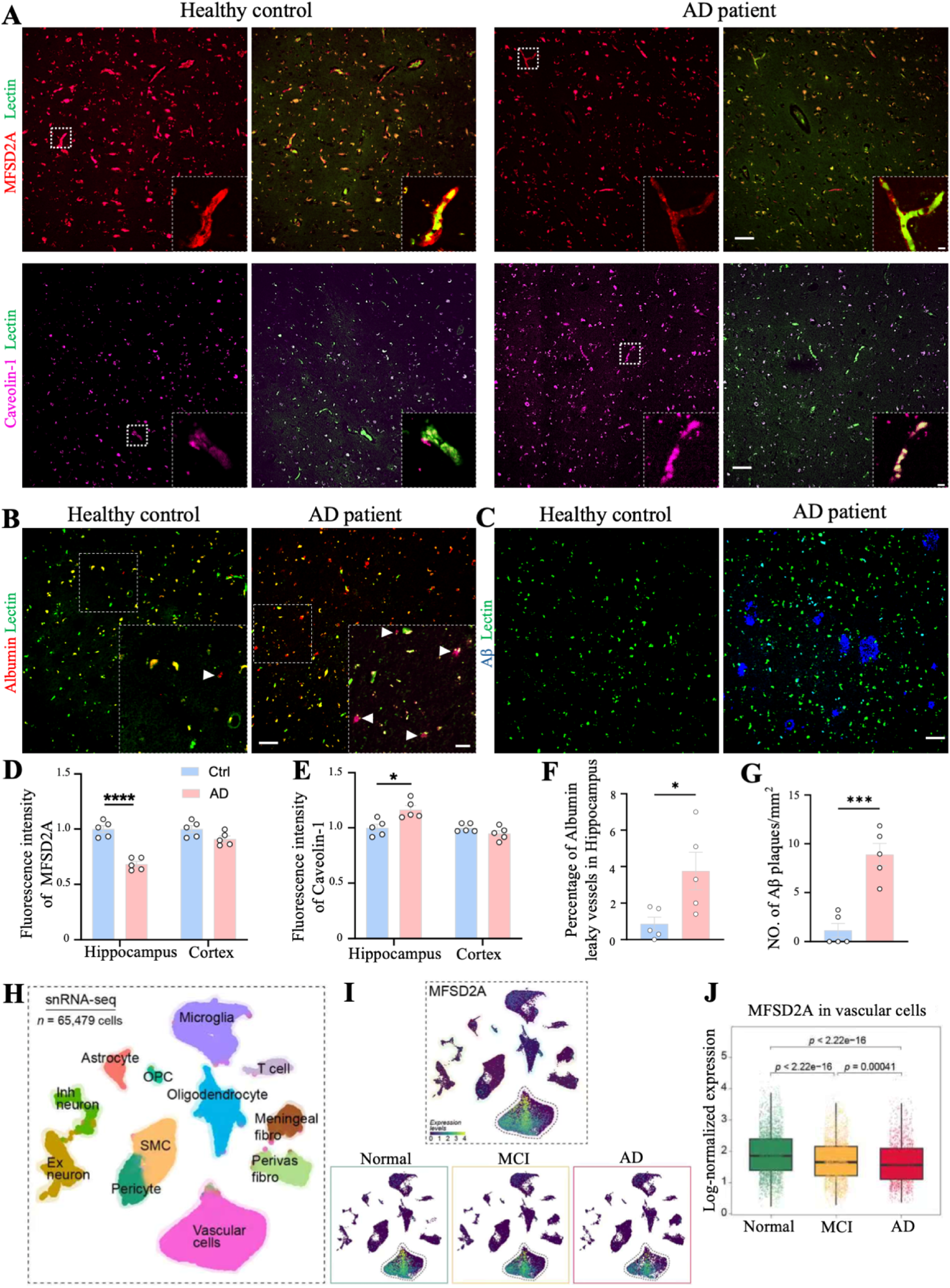
Reduced vascular MFSD2A and increased transcytosis in postmortem Alzheimer’s disease brains. (A) Representative immunofluorescence images of MFSD2A, caveolin-1, and lectin-labelled vessels in the hippocampus of control and AD brains. (B) Representative immunofluorescence images of hippocampal albumin extravasation. Arrowheads indicate perivascular albumin accumulation. (C) Representative immunofluorescence images of Aβ plaques and lectin-labelled vessels in the hippocampus of control and AD brains. (D) Quantification of vascular MFSD2A expression in the hippocampus and cortex. (E) Quantification of vascular caveolin-1 expression in the hippocampus and cortex. (F) Quantification of the percentage of albumin-leaky vessels in the hippocampus. (G) Quantification of hippocampal Aβ plaque density. (H–J) UMAP visualisation of 65,479 nuclei and MFSD2A expression in vascular cells from normal, mild cognitive impairment (MCI), and AD brains. Scale bars = 50 μm. Inset scale bars: 5 μm (A) and 20 μm (B). Data are mean ± SEM; *p < 0.05, ***p < 0.001, ****p < 0.0001; n = 5 donors/group; two-tailed unpaired Student’s *t*-test.

MFSD2A immunoreactivity was reduced by 31.6% ± 2.5% in the hippocampus of patients with AD compared with controls (Figure 1A, D). In contrast, MFSD2A levels in the cortex were only modestly affected (Supplementary Figure 1C; Figure 1D). Based on this, we next investigated whether this alteration was also evident at the transcriptomic level and detectable at earlier disease stages. We therefore interrogated a publicly available human single-nucleus RNA-sequencing dataset comprising 65,479 nuclei from individuals with normal cognition, mild cognitive impairment (MCI), and AD. Unsupervised clustering identified the major neuronal, neuroglial, immune, meningeal, and vascular cell populations (Figure 1H). *Mfsd2a* transcripts were predominantly confined to the vascular cell cluster (Figure 1I). The proportion of vascular cells decreased progressively along disease progression, from 37.1% in cognitively normal controls to 29.9% in MCI and 23.8% in AD (Supplementary Figure 1D). Within this population, MFSD2A expression was significantly reduced in MCI compared to cognitive normal controls and showed a further decline in AD (Figure 1J). These findings are consistent with our immunostaining results in AD brains and suggest that vascular MFSD2A decrease is present at MCI stage and becomes more pronounced in developed AD.

Caveolin-1, a key structural component of caveolae, is required for caveolar vesicle formation ^18^, while MFSD2A maintains BBB integrity in part by suppressing caveolae-mediated transcytosis ^11^. We next examined whether reduced vascular MFSD2A was accompanied by any alterations in caveolae-associated transcytosis. Albumin is a major circulating plasma protein which is normally excluded from the brain parenchyma. It may therefore serve as a functional readout of BBB permeability associated with endothelial transcytosis ^19^. Caveolin-1 immunoreactivity was significantly increased in the hippocampus of individuals with AD compared with controls (Figure 1A, E), whereas no significant difference was observed in the cortex (Supplementary Figure 1C; Figure 1E). Consistent with this finding, the proportion of hippocampal vessels exhibiting perivascular albumin extravasation was significantly higher in AD (Figure 1B, F). Together, these observations suggest that reduced vascular MFSD2A expression is associated with increased Caveolin-1 level and transcytosis-associated albumin extravasation in the human AD hippocampus, but not in the cortex.

### Hippocampus-selective MFSD2A reduction is associated with early BBB impairment in APP/PS1 mice

Based on the above observations in human samples, we next investigated whether MFSD2A levels undergo dynamic changes during AD progression in APP_SWE_/PS1_dE9_ (APP/PS1) mouse model. APP/PS1 mice exhibit elevated levels of soluble Aβ as early as 2-4 months of age, which subsequently progress to insoluble Aβ accumulation and extracellular plaque formation^20^. We assessed MFSD2A staining across different brain regions at 2, 4, and 10 months of age, representing distinct stages of disease progression. At 2 months of age (prior to disease onset), MFSD2A immunoreactivity was comparable between APP/PS1 and wild-type mice in all regions examined. By 4 months, however, MFSD2A levels were significantly reduced in the hippocampal dentate gyrus (DG) and Cornu Ammonis 1 (CA1) regions in APP/PS1 mice, whereas cortical levels remained similar between groups (Figure 2A, B-D). At 10 months of age, no significant genotype-dependent differences were detected in either hippocampal subregion or the cortex between the two mouse strains (Figure 2A, B-D). These results suggest an early and region-specific reduction in hippocampal MFSD2A protein levels during AD progression, as cortical and hippocampal MFSD2A levels were comparable in age-matched wild-type mice (Figure 2E). Together, these findings reveal an early stage and hippocampus-selective reduction in BBB MFSD2A protein level.

**Figure 2.**
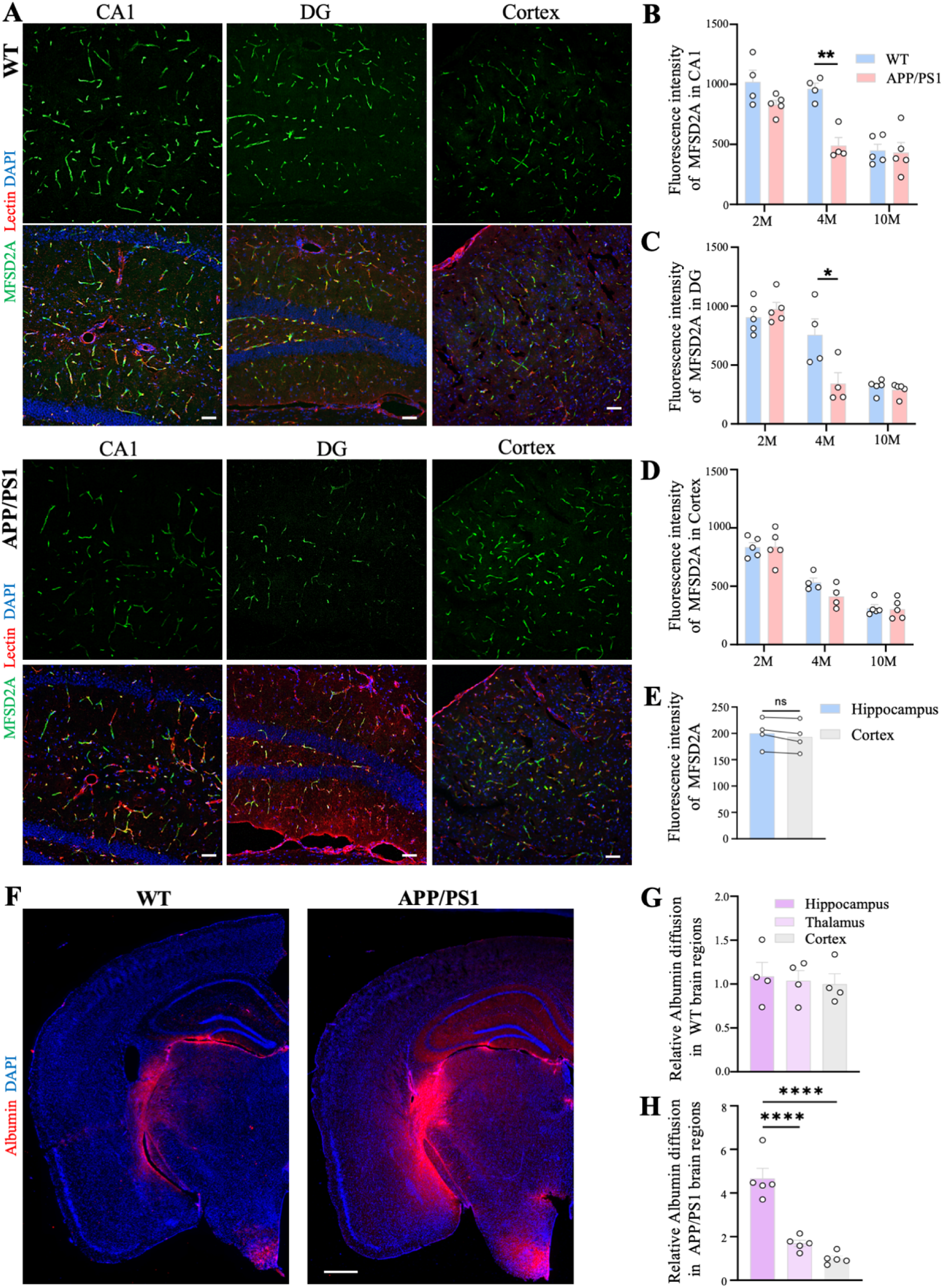
Early hippocampus-selective MFSD2A reduction and albumin leakage in APP/PS1 mice. (A) Representative immunofluorescence images of MFSD2A, lectin-labelled vessels, and DAPI in the CA1, DG, and cortex of 4-month-old WT and APP/PS1 mice. (B-D) Quantification of vascular MFSD2A expression in the CA1, DG, and cortex of 2-, 4-, and 10-month-old mice. (E) Comparison of MFSD2A expression between the two regions. (F) Representative immunofluorescence images of endogenous albumin distribution in WT and APP/PS1 brains. (G and H) Comparison of albumin diffusion among the hippocampus, thalamus, and cortex of WT and APP/PS1 mice. Scale bars, 50 μm (A) and 500 μm (F). Data are presented as mean ± SEM; each dot represents one mouse. Statistical significance was assessed using paired or unpaired two-tailed Student’s t-tests or one-way ANOVA. *p < 0.05, **p < 0.01, and ****p < 0.0001.

We next examined the regional distribution of parenchymal albumin accumulation in 5-month-old wild-type and APP/PS1 mice. In wild-type mice, albumin signal remained low in all BBB-protected brain regions and was primarily detected in areas with specialised fenestrated vasculature, such as the median eminence adjacent to the arcuate nucleus ^21,22^ (Figure 2F). In contrast, APP/PS1 mice showed a highly heterogeneous pattern of albumin accumulation across brain regions, with the hippocampus being the most affected (Figure 2F). Hippocampal albumin levels were 4.67 ± 0.46 times higher compared to those detected in the cortex and were also significantly higher than those in the thalamus (Figure 2G, H). This disproportionate hippocampal accumulation indicates a marked regional vulnerability of the BBB at early stage of amyloid pathology.

### MFSD2A overexpression at early amyloid stage mitigates hippocampal BBB transcytosis in APP/PS1 mice

To enable targeted upregulation of MFSD2A at the BBB, we generated an MFSD2A-expressing viral vector packaged in the AAV-BR1 capsid (AAV-BR1-MFSD2A), which preferentially transduces brain microvascular endothelial cells following systemic administration by injecting the tail vein ^23^ (Figure 3A). The corresponding control vector was designated AAV-BR1^CTL^. We first evaluated the efficiency of brain vascular targeting and tissue specificity of the AAV-BR1 vector using the encoded red fluorescent protein (RFP) as a reporter. Brain sections were co-stained for RFP and lectin to assess the spatial association between viral transduction and the cerebral vasculature, which showed that RFP staining was co-localised with lectin-positive cerebral blood vessels, with ∼62% of lectin-positive vessels exhibiting RFP immunoreactivity (Supplementary Figure 2A). To assess the peripheral biodistribution of the vector, RFP staining was further examined in the liver, spleen, lung, and kidney. No detectable RFP signal was observed in these organs (Supplementary Figure 2B). Collectively, these findings indicate that systemic administration of AAV-BR1 preferentially and efficiently transduces the brain microvascular endothelial cells, with negligible expression in major peripheral tissues.

**Figure 3.**
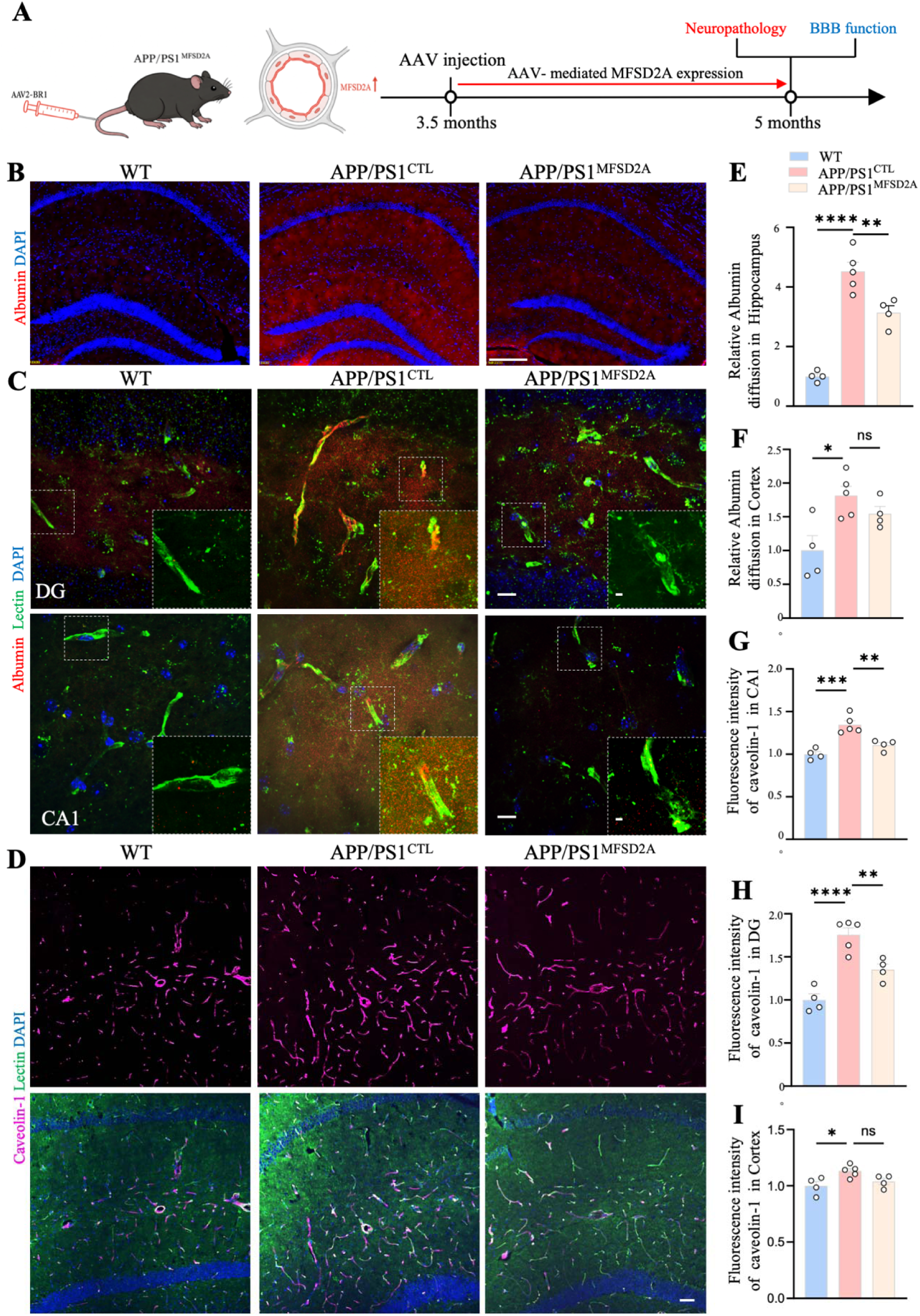
Brain endothelial MFSD2A overexpression suppresses caveolae-mediated transcytosis in early-stage APP/PS1 mice. (A) Schematic of AAV-BR1-mediated endothelial MFSD2A overexpression and the experimental timeline. Mice received AAV at 3.5 months of age and were analysed at 5 months. (B) Representative immunofluorescence images showing hippocampal albumin distribution in WT, APP/PS1^CTL^, and APP/PS1^MFSD2A^ mice. (C) Representative immunofluorescence images of albumin and lectin-labelled vessels in the DG and CA1. Insets show enlarged views of the boxed regions. (D) Representative immunofluorescence images of caveolin-1 and lectin-labelled vessels in the CA1. (E and F) Quantification of albumin extravasation in the hippocampus and cortex. (G–I) Quantification of vascular caveolin-1 fluorescence intensity in the CA1, DG, and cortex. Scale bars, 200 μm (B), 10 μm (C), and 50 μm (D). Inset scale bars, 2 μm (C). Data are presented as mean ± SEM; each dot represents one mouse. Statistical significance was assessed using one-way ANOVA. ns, not significant; *p < 0.05, **p < 0.01, ***p < 0.001, and ****p < 0.0001.

To determine whether overexpression of endothelial MFSD2A suppresses BBB transcytosis and prevents disease progression during the early disease stage, we specifically overexpressed endothelial MFSD2A in APP/PS1 mice (APP^MFSD2A^) at the age of 3.5 months. BBB function and neuropathological alterations were subsequently assessed at 5 months of age (Figure 3A). Albumin extravasation was evaluated as a marker of BBB permeability associated with endothelial transcytosis in the hippocampus and cortex. Compared to age-matched wild-type mice, APP/PS1^CTL^ mice exhibited a 4.53 ± 0.30-fold increase in hippocampal albumin extravasation (Figure 3B, E), which was partially attenuated by MFSD2A overexpression (Figure 3B, E). In contrast, cortical albumin extravasation was increased by only 1.81 ± 0.14-fold in APP/PS1^CTL^ mice, while there were no significant changes in APP/PS1^MFSD2A^ mice (Figure 3F). This indicates greater BBB vulnerability in the hippocampus during the early stage of AD, while endothelial MFSD2A overexpression limited vascular leakage in the hippocampus. To determine whether hippocampal albumin accumulation was spatially associated with local microvascular leakage, we examined albumin distribution relative to lectin-positive microvessels at higher magnification (Figure 3C), which demonstrated a marked increase in microvessel-associated albumin extravasation sites in 5-month-old APP/PS1^CTL^ mice compared to wild-type controls (Figure 3C). Notably, less perivascular albumin accumulation seen following MFSD2A overexpression, is consistent with the quantitative analysis, suggesting that endothelial MFSD2A overexpression limits albumin leakage across the hippocampal BBB. Because MFSD2A inhibits endothelial transcytosis by suppressing caveolae formation^11^, we next examined caveolin-1, a key structural component of caveolae^18^, to determine whether the reduction in albumin leakage was accompanied by altered caveolar transport pathway. In the hippocampus, caveolin-1 immunoreactivity was markedly increased in both CA1 and DG of 5-month-old APP/PS1^CTL^ mice (Figure 3D, G, H; Supplementary Figure 2C); immunoreactivity of caveolin-1 was significantly attenuated by MFSD2A overexpression (Figure 3 D, G, H; Supplementary Figure 2C). In the cortex, caveolin-1 levels were modestly increased in APP/PS1 mice, and were not affected by MFSD2A overexpression (Figure 3I; Supplementary Figure 2D), indicating transcytosis-driven albumin leakage in the hippocampal BBB.

### Early stage MFSD2A overexpression ameliorates hippocampal BBB dysfunction and reduces amyloid pathology in APP/PS1 mice

We next examined two key BBB-associated proteins known to be altered in AD, as previously shown by us and others, namely glucose transporter 1 (GLUT1) and tight junction protein Claudin-5 ^3,24,25^. GLUT1 is the dominant insulin-independent glucose transporter at the BBB, which is consistently reduced in both AD patients and murine models of AD ^3,24,26^. In line with previous observations, in 5-month-old APP/PS1^CTL^ mice, GLUT1 immunoreactivity was significantly lower in the hippocampal CA1 (Figure 4A, B) and DG regions (Figure 4C; Supplementary Figure 3A), as well as in the cortex (Figure 4D; Supplementary Figure 3A, B). APP/PS1^MFSD2A^ showed normalised GLUT1 levels in both hippocampal CA1 and DG, with unchanged cortical GLUT1 levels (Figure 4A-D; Supplementary Figure 3A, B). Claudin-5 loss has also been implicated in BBB impairment in AD progression, including hippocampal barrier malfunction in early stage of Aβ pathology in APP/PS1 mice^3^. Here, Claudin-5 immunoreactivity was significantly reduced in the hippocampus of 5-month-old APP/PS1^CTL^ mice (Figure 4E, F), which was rescued in APP/PS1^MFSD2A^ (Figure 4E, F). These findings suggest that endothelial MFSD2A overexpression can normalise the levels of BBB-associated glucose transporter and junctional proteins in the hippocampus during early stage of AD pathology.

**Figure 4.**
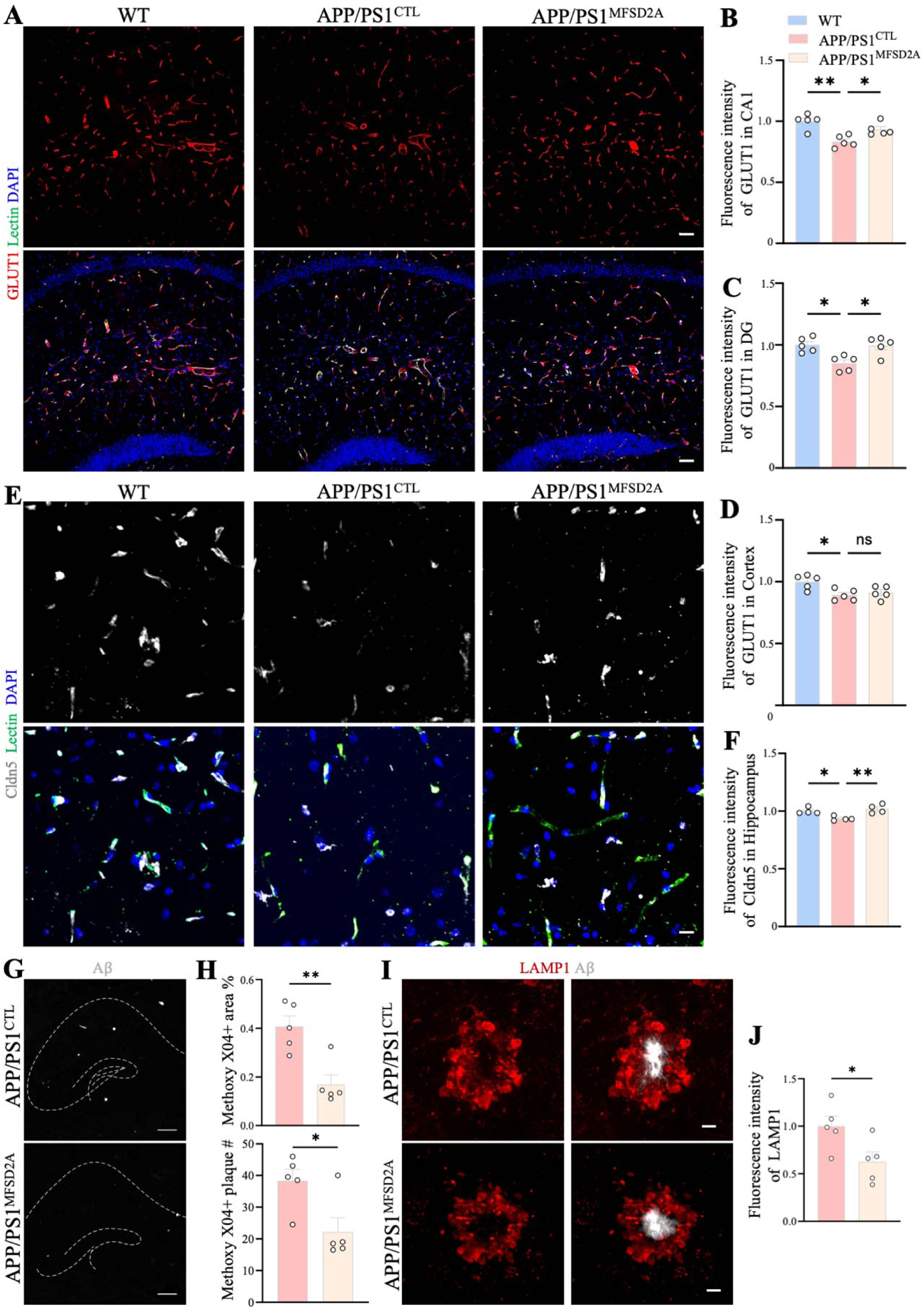
MFSD2A overexpression at early amyloid stages alleviates BBB dysfunction and neurodegeneration in APP/PS1 mice. (A) Representative immunofluorescence images of GLUT1 and lectin-labelled vessels in the CA1 of WT, APP/PS1^CTL^, and APP/PS1^MFSD2A^ mice. (B–D) Quantification of vascular GLUT1 fluorescence intensity in the CA1, DG, and cortex. (E) Representative immunofluorescence images of claudin-5 and lectin-labelled vessels in the hippocampus. (F) Quantification of vascular claudin-5 fluorescence intensity in the hippocampus. (G) Representative images of Methoxy-X04-labelled Aβ plaques in the hippocampus of APP/PS1^CTL^ and APP/PS1^MFSD2A^ mice. Dashed lines delineate the hippocampal region. (H) Quantification of Aβ plaque number and plaque burden. (I) Representative immunofluorescence images of LAMP1 immunoreactivity surrounding Aβ plaques. (J) Quantification of plaque-associated LAMP1 fluorescence intensity. Scale bars, 50 μm (A), 20 μm (E), 200 μm (G), and 5 μm (I). Data are presented as mean ± SEM; each dot represents one mouse. Statistical significance was assessed using one-way ANOVA or an unpaired two-tailed Student’s t-test. ns, not significant; *p < 0.05 and **p < 0.01.

We next assessed AD neuropathology following endothelial MFSD2A overexpression. We found significant reductions in both Aβ plaque burden and plaque number in APP/PS1^MFSD2A^ mice compared with APP/PS1^CTL^ mice (Figure 4G, H). We further examined lysosomal-associated membrane protein 1 (LAMP1), a marker of dystrophic neurites surrounding amyloid plaques ^27^. Consistent with the amelioration of Aβ pathology, immunoreactivity of LAMP1 was also significantly decreased following MFSD2A overexpression (Figure 4I, J). Together, these findings suggest that endothelial MFSD2A overexpression can reduce Aβ deposition and attenuate neuritic dystrophy in APP/PS1 mice.

### Late-stage endothelial MFSD2A overexpression is insufficient to reverse established BBB dysfunction and cognitive impairment

To determine whether MFSD2A overexpression is effective at the advanced stage of AD-associated pathology with pronounced cognitive deficits, we performed a late-stage intervention, where APP/PS1 mice received a single tail-vein injection of AAV-BR1-MFSD2A at 9 months of age and underwent behavioural testing at 10 months, followed by histological assessment of BBB alterations and neuropathology (Figure 5A).

**Figure 5.**
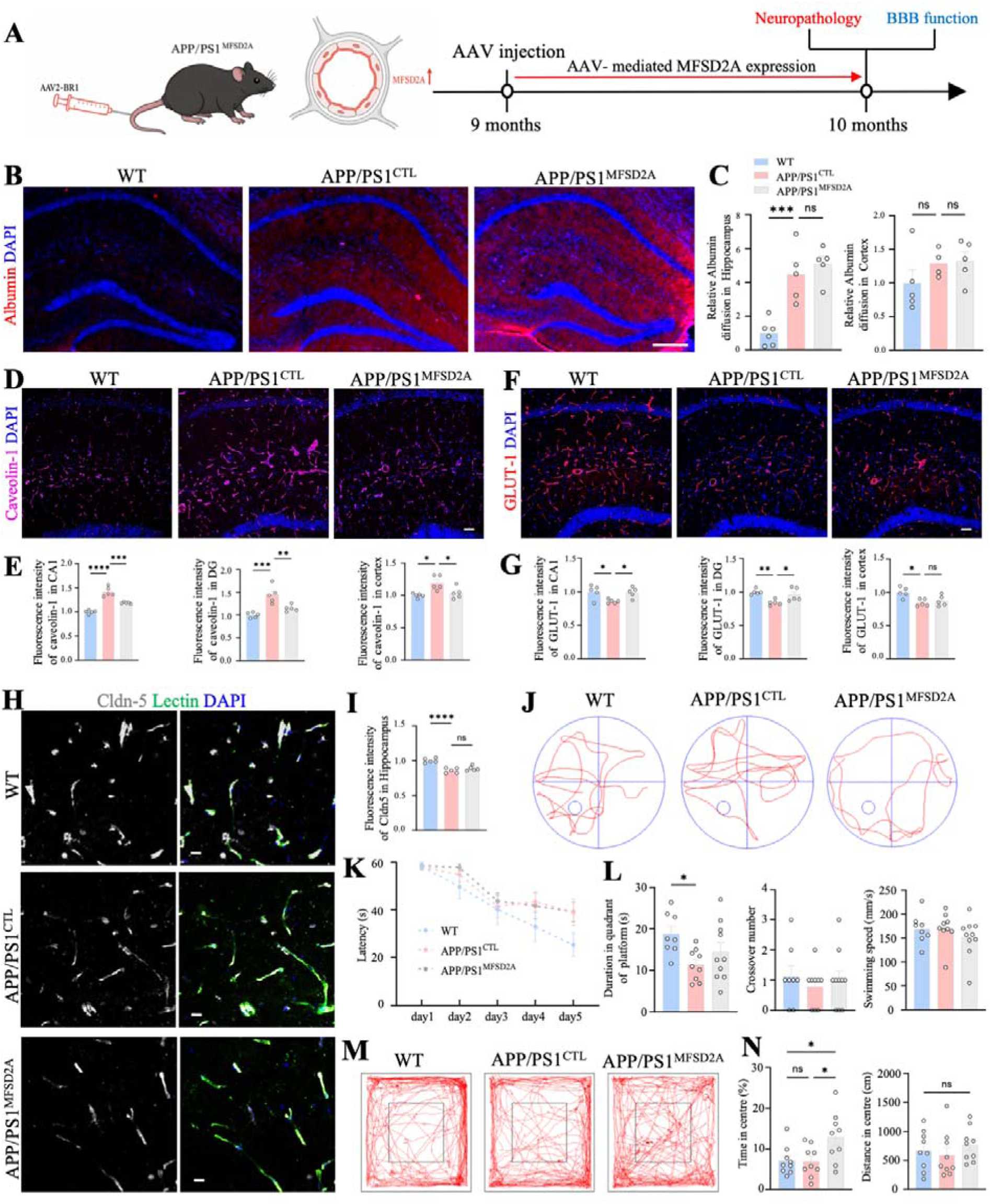
MFSD2A overexpression at later stage of the disease does not alleviate BBB malfunction and neurological outcome in APP/PS1 mice. (A) Schematic of late-stage AAV-BR1-mediated endothelial MFSD2A overexpression and the experimental timeline. Mice received AAV at 9 months of age and were analysed at 10 months. (B) Representative immunofluorescence images showing hippocampal albumin distribution in WT, APP/PS1^CTL^, and APP/PS1^MFSD2A^ mice. (C) Quantification of albumin extravasation in the hippocampus and cortex. (D) Representative immunofluorescence images of vascular caveolin-1 expression in the CA1. (E) Quantification of vascular caveolin-1 fluorescence intensity in the CA1, DG, and cortex. (F) Representative immunofluorescence images of vascular GLUT1 expression in the CA1. (G) Quantification of vascular GLUT1 fluorescence intensity in the CA1, DG, and cortex. (H) Representative immunofluorescence images of claudin-5 and lectin-labelled vessels in the hippocampus. (I) Quantification of vascular claudin-5 fluorescence intensity in the hippocampus. (J) Representative swimming trajectories during the Morris water maze probe trial. (K) Escape latency during the 5-day acquisition phase. (L) Time spent in the target quadrant, number of platform crossings, and swimming speed during the probe trial. (M) Representative trajectories in the open-field test. (N) Quantification of time spent and distance travelled in the centre area. Scale bars, 200 μm (B), 50 μm (D and F), and 20 μm (H). Data are presented as mean ± SEM; each dot represents one mouse. Statistical significance was assessed using one-way ANOVA. ns, not significant; *p < 0.05, **p < 0.01, ***p < 0.001, and ****p < 0.0001.

Hippocampal albumin extravasation was significantly increased in 10-month-old APP/PS1^CTL^ mice compared with age-matched wild-type controls (Figure 5B, C). In contrast to the marked reduction observed following early-stage intervention, late-stage MFSD2A overexpression did not alleviate hippocampal albumin accumulation (Figure 5B, C). No significant difference in cortical albumin extravasation was observed among the three groups (Figure 5C). Caveolin-1 levels were significantly increased in both the CA1 and DG of APP/PS1^CTL^ mice and were attenuated in APP/PS1^MFSD2A^ in both hippocampal subregions (Figure 5D, E), albeit with no changes in extravascular albumin accumulation. GLUT1 levels, reduced in the CA1 and DG of 10-month-old APP/PS1 mice, were restored following MFSD2A overexpression, whereas cortical GLUT1 levels remained unchanged (Figure 5F, G). Consistent with BBB disruption during disease progression, hippocampal Claudin-5 levels were significantly decreased in 10-month-old APP/PS1^CTL^ mice and were unaffected by MFSD2A overexpression (Figure 5H, I). These findings suggest that MFSD2A overexpression at an advanced AD stage normalises some BBB functional markers, but is insufficient to reverse established albumin accumulation and tight junctional disruption.

Exploratory behaviour and general locomotor activity were subsequently assessed using the open-field test (Figure 5M). Time spent in the centre did not differ significantly between wild-type and APP/PS1^CTL^ mice (Figure 5N). Of note, APP/PS1^MFSD2A^ mice spent significantly more time in the central area than both wild-type and APP/PS1^CTL^ mice (Figure 5N; Supplementary Figure 4D-F). Spatial learning and memory were subsequently evaluated using the Morris water maze (Figure 5J). During the acquisition phase, APP/PS1^CTL^ mice exhibited significantly increased escape latency compared with wild-type controls, whereas APP/PS1^MFSD2A^ mice displayed a learning curve comparable to APP/PS1^CTL^ mice (Figure 5K). In the probe trial, APP/PS1^CTL^ mice spent significantly less time in the target quadrant than wild-type controls, consistent with impaired spatial memory (Figure 5L). This impairment was not rescued by late-stage MFSD2A overexpression (Figure 5L). The number of platform crossings and swimming velocity did not differ significantly among groups (Figure 5L).

We next examined whether late-stage MFSD2A overexpression altered established amyloid pathology. There were no significant differences in hippocampal Aβ plaque number or plaque burden between APP/PS1^MFSD2A^ and APP/PS1^CTL^ mice (Supplementary Figure 4A, B). Similarly, LAMP1 immunoreactivity around Aβ plaques remained unchanged following MFSD2A overexpression (Supplementary Figure 4C). These findings suggest that late-stage MFSD2A overexpression is insufficient to attenuate established amyloid and plaque-associated neuritic pathology.

Collectively, these findings demonstrate that endothelial MFSD2A overexpression initiated at an advanced stage of Aβ pathology in APP/PS1 mice can mitigate abnormal hippocampal caveolin-1 and GLUT1 expression but is insufficient to restore BBB integrity or cognitive function when AD pathological alterations are established.

### Early and sustained MFSD2A overexpression preserves BBB function in APP/PS1 mice

Early-stage MFSD2A overexpression ameliorated BBB malfunction when assessed at 5 months, whereas intervention initiated at 9 months showed limited efficacy against established BBB damage. These findings raised the possibility that long-term protection requires MFSD2A restoration before advanced barrier disruption and continued intervention during disease progression. To test this hypothesis, APP/PS1 mice received MFSD2A-expressing or control vector at 3.5 months of age, and a second injection of the corresponding vector was administered at 6.5 months (termed as APP/PS1^S-MFSD2A^ and APP/PS1^S-CTL^ mice, respectively) to sustain MFSD2A expression (Figure 6A).

**Figure 6.**
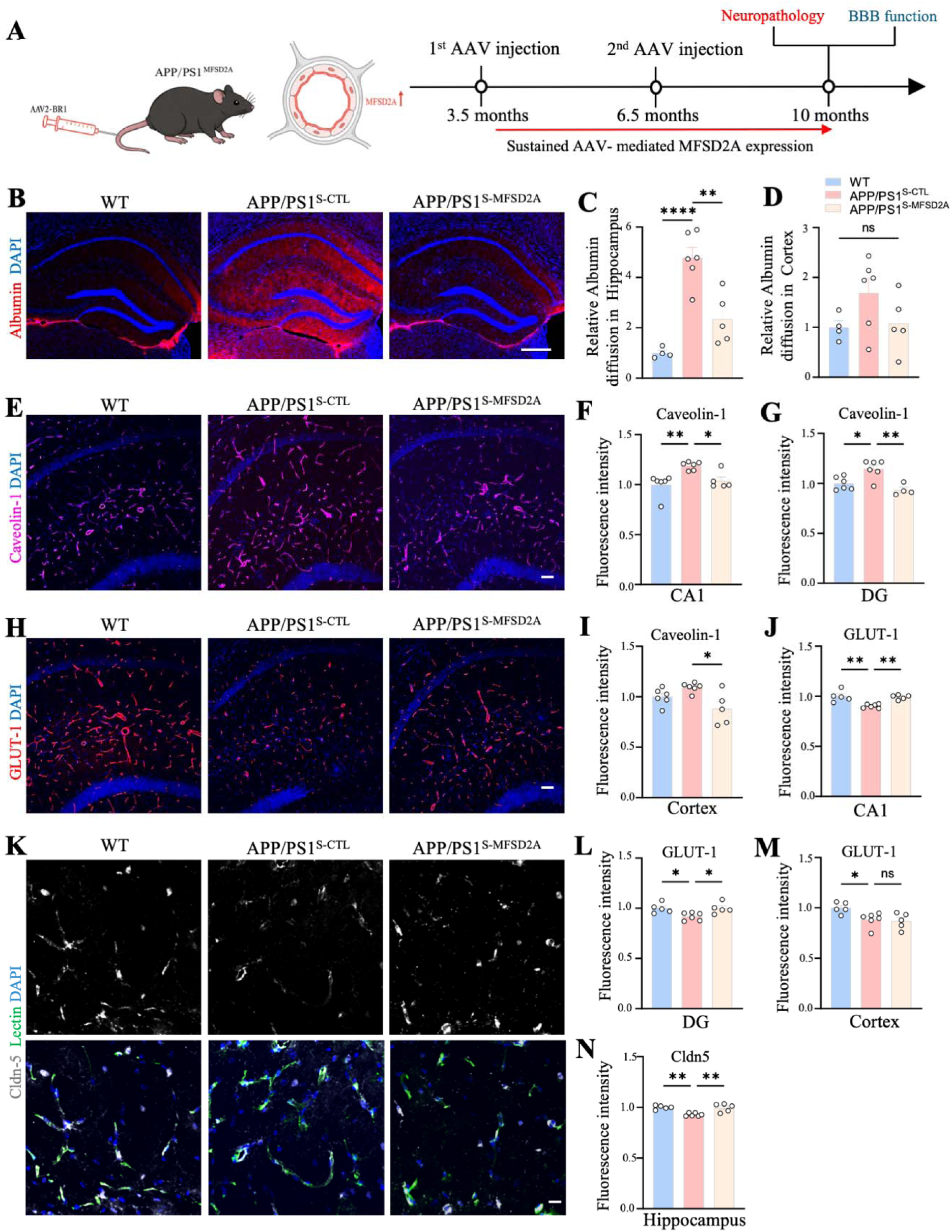
Early and sustained MFSD2A overexpression rescues BBB dysfunction in APP/PS1 mice. (A) Schematic of early and sustained AAV-BR1-mediated endothelial MFSD2A overexpression and the experimental timeline. Mice received AAV injections at 3.5 and 6.5 months of age and were analysed at 10 months. (B) Representative immunofluorescence images showing hippocampal albumin distribution in WT, APP/PS1^S-CTL^, and APP/PS1^S-MFSD2A^ mice. (C and D) Quantification of albumin extravasation in the hippocampus and cortex. (E) Representative immunofluorescence images of vascular caveolin-1 expression in the CA1. (F, G, and I) Quantification of vascular caveolin-1 fluorescence intensity in the CA1, DG, and cortex. (H) Representative immunofluorescence images of vascular GLUT1 expression in the CA1. (J, L, and M) Quantification of vascular GLUT1 fluorescence intensity in the CA1, DG, and cortex. (K) Representative immunofluorescence images of claudin-5 and lectin-labelled vessels in the hippocampus. (N) Quantification of vascular claudin-5 fluorescence intensity in the hippocampus. Scale bars, 200 μm (B), 50 μm (E and H), and 20 μm (K). Data are presented as mean ± SEM; each dot represents one mouse. Statistical significance was assessed using one-way ANOVA. ns, not significant; *p < 0.05, **p < 0.01, and ****p < 0.0001.

Immunofluorescence staining showed substantial parenchymal albumin accumulation in the hippocampus of APP/PS1^S-CTL^ mice. This accumulation was significantly reduced in APP/PS1^S-MFSD2A^ mice, as confirmed by quantitative analysis of hippocampal albumin immunoreactivity (Figure 6B, C). Consistent with this reduction, hippocampal caveolin-1 immunoreactivity was also significantly lower in APP/PS1^S-MFSD2A^ mice than in APP/PS1^S-CTL^ mice (Figure 6E-G; Supplementary Figure 5A). These findings suggest that early and sustained MFSD2A overexpression can suppress caveolin-1 levels and limit transcytosis-associated albumin accumulation during AD progression.

We next examined immunoreactivity for GLUT1 and Claudin-5. GLUT1 expression was reduced in the hippocampus of 10-month-old APP/PS1^S-CTL^ mice, which was normalised by sustained MFSD2A overexpression (Figure 6H, J, L; Supplementary Figure 5B). Early and prolonged MFSD2A overexpression also prevented the decline in hippocampal Claudin-5 levels (Figure 6K, N). Together, these results show that early and sustained MFSD2A overexpression limits hippocampal albumin accumulation, preserves multiple BBB-associated markers, including GLUT1 and Claudin-5, thereby providing more durable protection against BBB dysfunction than intervention initiated at a later disease stage.

### Early and sustained endothelial MFSD2A overexpression attenuates amyloid pathology and improves hippocampal function in APP/PS1 mice

We then assessed whether early and sustained MFSD2A overexpression altered neuropathological progression in APP/PS1 mice. At 10 months of age, both Aβ plaque number and area were significantly reduced in APP/PS1^S-MFSD2A^ compared with APP/PS1^S-CTL^ (Figure 7A-C). These changes were accompanied with marked reductions in plaque-associated LAMP1 and Reticulon-3 (RTN3) immunoreactivity (Figure 7D-G), suggesting attenuated neuritic dystrophy. To determine whether the improvements in BBB function and neuropathology were accompanied by functional preservation of hippocampal circuits, we examined long-term potentiation in acute hippocampal slices (Figure 7H). APP/PS1 ^S-CTL^ mice exhibited markedly impaired LTP compared with wild-type mice, as indicated by a sustained reduction in the normalised fEPSP slope following LTP induction (Figure 7I). Early and sustained MFSD2A overexpression significantly enhanced LTP in APP/PS1 mice, with an increased fEPSP slope during the late phase of the recording compared to sham-treated APP/PS1 mice (Figure 7I), suggesting improved hippocampal synaptic plasticity in APP/PS1 mice.

**Figure 7.**
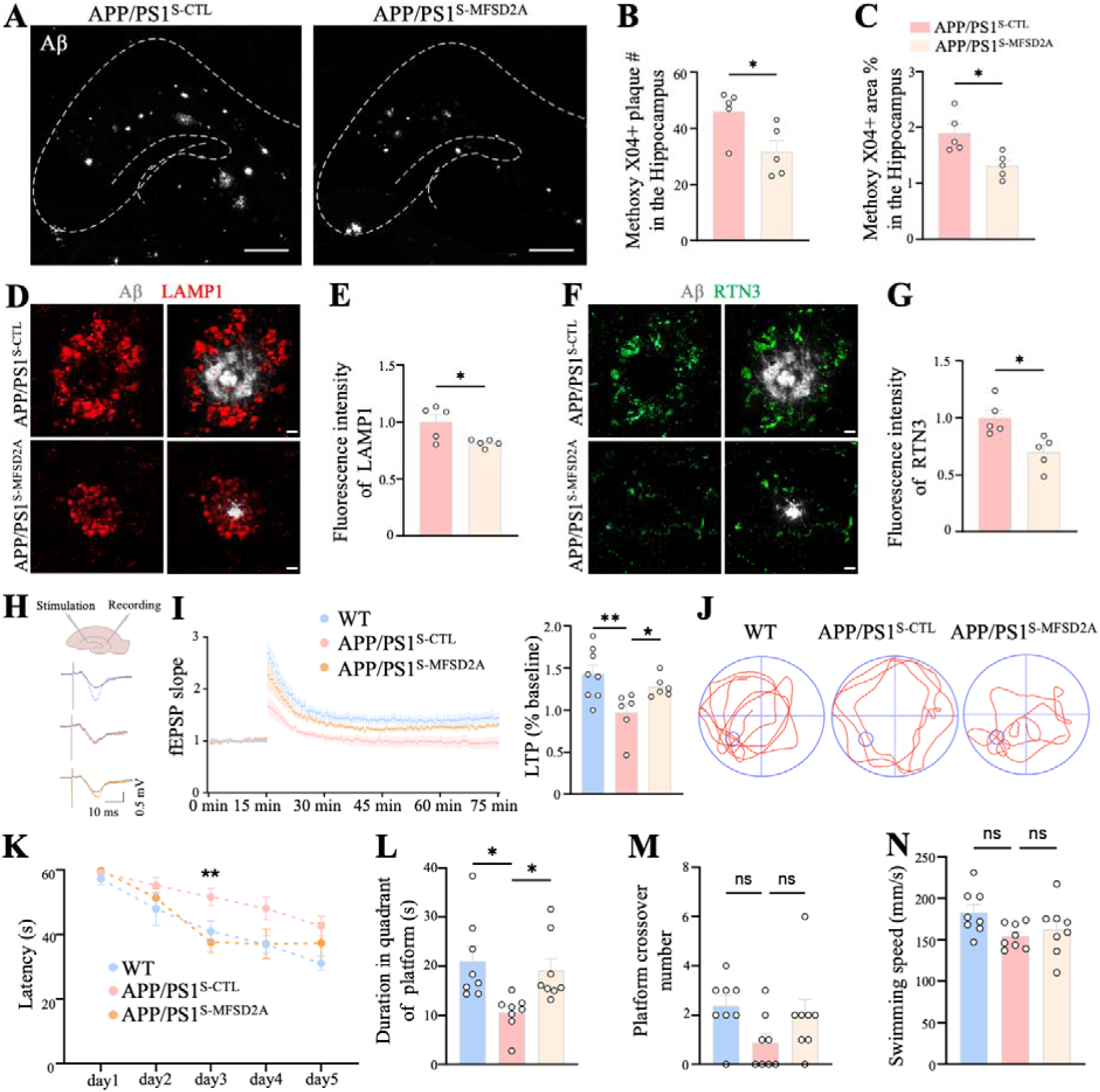
Early and sustained MFSD2A overexpression improves neurological outcome in APP/PS1 mice. (A) Representative images of Methoxy-X04-labelled Aβ plaques in the hippocampus of APP/PS1^S-CTL^ and APP/PS1^S-MFSD2A^ mice. Dashed lines delineate the hippocampal region. (B and C) Quantification of hippocampal Aβ plaque number and plaque burden. (D) Representative immunofluorescence images of LAMP1 immunoreactivity surrounding Aβ plaques. (E) Quantification of plaque-associated LAMP1 fluorescence intensity. (F) Representative immunofluorescence images of RTN3 immunoreactivity surrounding Aβ plaques. (G) Quantification of plaque-associated RTN3 fluorescence intensity. (H) Schematic of hippocampal LTP recording with representative field excitatory postsynaptic potential (fEPSP) traces. (I) Time course and quantification of long-term potentiation, expressed as the normalized fEPSP slope relative to baseline. (J) Representative swimming trajectories during the Morris water maze probe trial. (K) Escape latency during the 5-day acquisition phase. (L–N) Time spent in the target quadrant, number of platform crossings, and swimming speed during the probe trial. Scale bars, 200 μm (A), 5 μm (D), and 5 μm (F). Data are presented as mean ± SEM. Statistical significance was assessed using an unpaired two-tailed Student’s t-test, one-way ANOVA. ns, not significant; *p < 0.05 and **p < 0.01.

We next investigated whether early and sustained MFSD2A overexpression could preserve cognitive function. Spatial learning and memory were assessed in the Morris water maze at 10 months (Figure 7J). During the acquisition phase, APP/PS1^S-CTL^ mice exhibited prolonged escape latencies compared with age-matched wild-type controls, whereas APP/PS1^S-MFSD2A^ showed mice significantly shortened escape latency than APP/PS1^S-CTL^ mice (Figure 7K). In the probe trial, APP/PS1^S-CTL^ mice spent significantly less time in the target quadrant than wild-type controls, and this deficit was significantly ameliorated by sustained MFSD2A overexpression (Figure 7L). The number of crossings over the former platform location and swimming velocity did not differ significantly among the groups (Figure 7M, N). Together, these findings suggest that the broad protective effects of sustained MFSD2A overexpression extended beyond BBB and neuropathological outcomes to include improved spatial learning and memory in AD mice.

## Discussion

BBB malfunction is increasingly recognised as an early contributor to AD, yet the endothelial mechanisms governing regional vulnerability and therapeutic responsiveness remained largely unknown. Here, we identified endothelial MFSD2A loss as an early molecular signature of impaired hippocampal BBB in both human AD and APP/PS1 mice. We demonstrate that restoration of MFSD2A can preserve BBB integrity and modify disease progression when delivered during early pathology. Our study establishes the MFSD2A-caveolae pathway as a stage-dependent and therapeutically accessible regulator of neurovascular dysfunction in AD (Figure 8).

**Figure 8.**
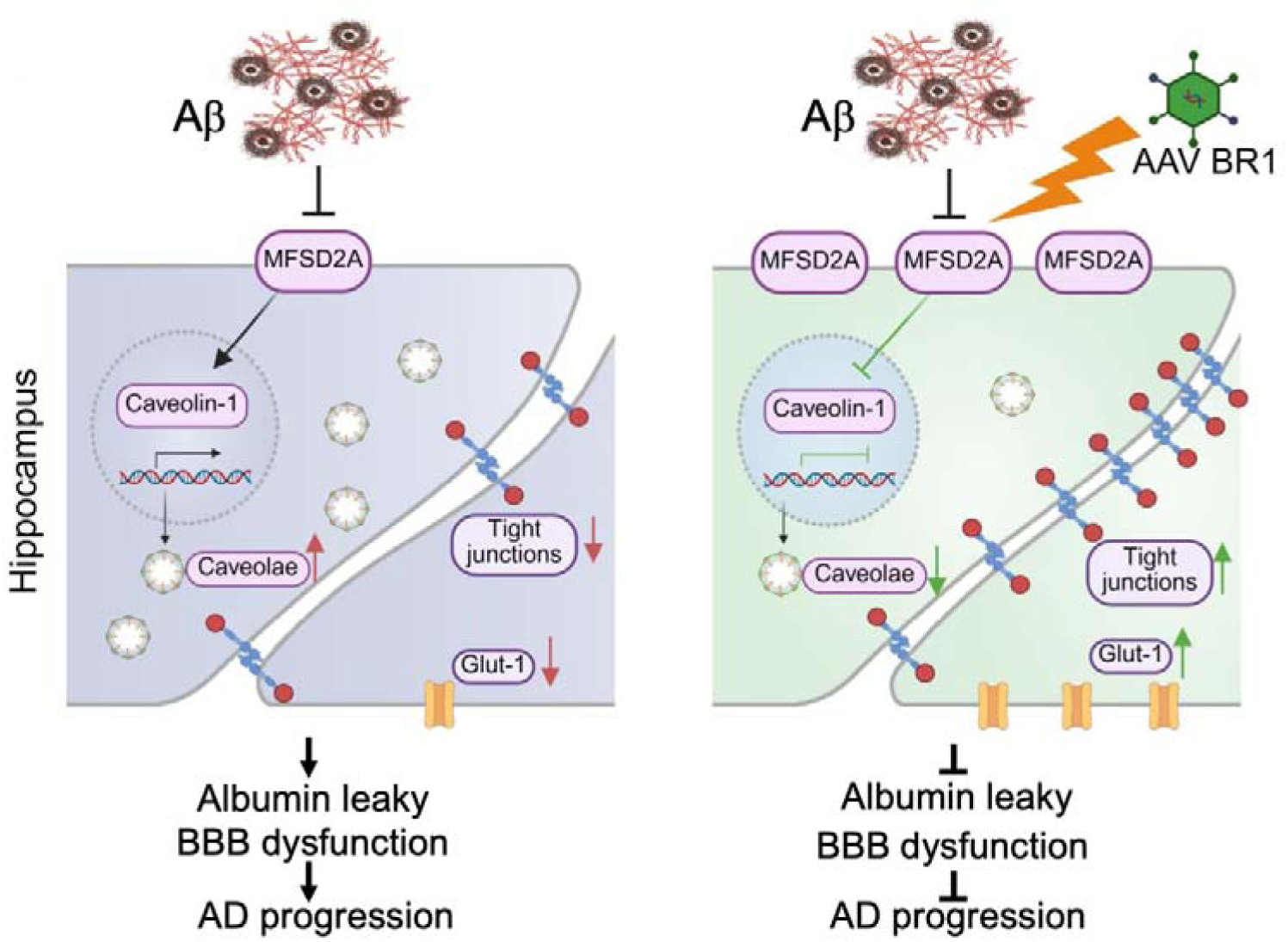
Proposed model of early endothelial MFSD2A protection against hippocampal BBB dysfunction in AD. Left: Early hippocampal MFSD2A reduction is associated with increased caveolin-1 expression, enhanced caveolae-mediated transcytosis, albumin leakage, and reduced GLUT1 and claudin-5 levels. Right: Early and sustained AAV-BR1-mediated endothelial MFSD2A overexpression reduces caveolin-1 expression and albumin accumulation, preserves GLUT1 and claudin-5, and attenuates amyloid-associated pathology and hippocampal dysfunction.

Key findings of this study are identification of marked regional heterogeneity of BBB impairment and its association with endothelial MFSD2A loss. Although amyloid pathology is present in both cortex and hippocampus of APP/PS1 mice, BBB disruption, reflected by albumin accumulation, was disproportionately higher in the hippocampus where MFSD2A reduction was most pronounced. This regional difference was not explained by intrinsically lower MFSD2A levels in the hippocampus, as hippocampal and cortical vessels exhibited comparable baseline MFSD2A levels in wild-type mice. These observations suggest that the vascular response to amyloid pathology is not determined solely by regional plaque burden. Rather, the hippocampal BBB appears particularly susceptible to the loss of the MFSD2A-dependent transcellular barrier. The upstream mechanisms of MFSD2A loss in AD remain unclear. Previous studies suggest that MFSD2A is positively regulated by Wnt/β-catenin signalling^28^, whereas TGF-β1 suppresses its transcription through Tgfbr2-Smad2/4 signalling, thereby promoting caveolae-mediated transcytosis^29^. Whether these pathways account for the early and region-selective loss of MFSD2A observed in the hippocampus requires further investigation. Recent studies have highlighted substantial regional specialisation of the BBB, including endothelial heterogeneity^17,30,31^. Our findings extend this concept by identifying MFSD2A as a potential molecular mechanism underlying hippocampal BBB vulnerability to amyloid pathology in AD. Such BBB heterogeneity may help explain the dissociation between the widespread anatomical distribution of amyloid pathology and the preferential impairment of hippocampus-dependent memory. Although MFSD2A loss is unlikely to be the sole driver of this susceptibility, our findings suggest endothelial regulation of transcytosis as a previously underappreciated contributor to region-dependent neuropathology during AD progression.

Caveolin-1 and perivascular albumin accumulation changes in parallel to changes in MFSD2A, are consistent with dysregulation of caveolae-associated transcytosis. MFSD2A defines the low rate of non-selective vesicular transport that characterises the BBB ^10,11^. In this context, the increase in Caveolin-1 and perivascular albumin extravasation is most readily interpreted as loss of a physiological suppression of transcellular barrier function. The transcytosis is not inherently detrimental. Receptor- and carrier-mediated transport is essential for nutrient and hormone delivery, intercellular communications, and the clearance of brain-derived metabolites ^12,32,33^. The pathological feature is therefore not transcytosis *per se*, but the loss of transport selectivity that permits excessive entry of circulating macromolecules into the brain parenchyma. Albumin leak is of particular interest because extravasation can activate astrocytic TGFβ signalling, disrupt neuronal networks and prompt cognitive impairment ^34^. Restoring MFSD2A is therefore not intended to abolish endothelial transport, but rather to re-establish the highly selective transport state that defines a functional BBB while preserving physiological exchange pathways.

The distinct effects of early, late, and sustained intervention indicate that therapeutic efficacy of MFSD2A restoration strongly depends on the disease stage. Early intervention improved Caveolin-1, GLUT1, and Claudin-5 and limited albumin leakage, whereas late intervention although normalising Caveolin-1 and GLUT1, failed to reduce albumin extravasation or restore Claudin-5. These findings suggest that different components of BBB pathology differ in their reversibility. Endothelial transcytosis and transporter-associated phenotypes may remain relatively plastic after amyloid pathology is established, whereas junctional abnormalities and accumulated parenchymal injury seems to be less responsive. Although increased transcytosis has been shown to precede tight-junction disruption in acute stroke models ^13^, the temporal relationship between these barrier pathways is likely to be disease- and model-dependent. In our study, Claudin-5 was already reduced at the early stage of pathology ^3^, suggesting that transcellular and paracellular barrier defects may arise concurrently and potentially reinforce one another ^35^. The superior outcome of sustained MFSD2A restoration further suggests that continued endothelial protection may be required as amyloid pathology progresses.

The late-intervention cohort revealed an apparent dissociation between Caveolin-1 and albumin. Although MFSD2A restoration reduced Caveolin-1 abundance, hippocampal albumin accumulation remained unchanged. This dissociation may reflect the multifactorial nature of BBB malfunction at advanced disease stages. Once tight-junction integrity is compromised, albumin entry may continue through paracellular routes despite suppression of Caveolin-1-dependent transcytosis ^13^. Moreover, AD-associated impairment of interstitial and meningeal lymphatic drainage may delay the removal of extravasated albumin ^36^. Consequently, tissue albumin levels may reflect both ongoing leakage and the persistence of previously accumulated protein. Together, continued Caveolin-1-independent entry and impaired clearance may sustain parenchymal albumin accumulation despite partial restoration of endothelial transcytosis. This interpretation is supported by the sustained-intervention experiment, in which MFSD2A overexpression was initiated before advanced pathology and maintained throughout disease progression, resulting in substantially lower hippocampal albumin accumulation. These findings suggest that preventing continued plasma-protein entry may be more achievable than removing an established parenchymal protein burden.

The sustained reduction in Aβ plaque burden, plaque number, and dystrophic neurites following MFSD2A overexpression further indicates that preservation of the BBB favours broader neuropathological benefits.

Restricting entry of albumin and other plasma constituents may reduce secondary glial and synaptic injury ^34,37,38^. Preservation of GLUT1 may support cerebral energy supply and metabolism ^39,40^, whereas improved BBB homeostasis may facilitate physiological Aβ clearance ^41^. Consistent with these neuropathological improvements, sustained MFSD2A restoration enhanced hippocampal long-term potentiation and improved performance in the Morris water maze. Conversely, late intervention failed to improve spatial learning and memory despite normalising selected endothelial markers, indicating that partial restoration of endothelial function is insufficient once downstream neural dysfunction is established.

Several limitations should be considered. Human sample analyses were cross-sectional due to the innate limitation. In addition, although AAV-BR1 preferentially targeted brain microvascular endothelium, it transduced only a subset of cerebral vessels, and vector-driven MFSD2A expression may not fully recapitulate native MFSD2A expression and vascular distribution. Finally, the APP/PS1 mouse model recapitulates amyloid pathology but does not capture the full complexity of sporadic AD, including tau pathology, vascular comorbidities and other age-related contributors to disease progression ^42,43^.

In summary, our findings identify endothelial MFSD2A loss as an early and region-specific feature of AD-associated BBB dysfunction. They further demonstrate that the consequences of MFSD2A restoration are highly dependent on timing of therapy, with early and sustained intervention preserving BBB integrity, limiting neuropathology, maintaining synaptic plasticity, and improving cognitive function. Conversely, late intervention modified selected endothelial markers but failed to reverse established vascular and neural dysfunction. These findings position the MFSD2A–caveolae axis as a therapeutically targetable regulator of neurovascular dysfunction and suggest that vascular intervention is most effective before BBB breakdown coupled to persistent protein accumulation, junctional injury, and neural dysfunction.

## Data availability

All data that support the findings of this study are available from the corresponding author upon reasonable request.

## Acknowledgements

This work was supported by National Natural Science Foundation of China (82501901 to Y.Z.), Shenzhen Fundamental Research Program (RCJC20231211090018040 and ZDSYS20220606100801003 to C.Y.; JCYJ20250604143752069 to Y.Z.), and “3510” Medical Innovation Leadership Program, Fundamental Research Funds for the Central Universities, Sun Yat-sen University (20263510JCQN011 to C.Y.). Human tissue was provided by the National Human Brain Bank for Development and Function, Chinese Academy of Medical Sciences and Peking Union Medical College, Beijing, China. This study was supported by the Institute of Basic Medical Sciences, Chinese Academy of Medical Sciences, Neuroscience Centre, and the China Human Brain Banking Consortium, by grants from STI2030-Major Project #2021ZD0201100 Task 1 #2021ZD0201101. We thank Lan Yuan from the Biomedical lmaging Core Facility at the Bio-Tech Centre of Shenzhen Medical Academy of Research and Translation (SMART) for the SpinSR spinning disk confocal microscopy technical support.

---

**Supplementary Figure 1.**
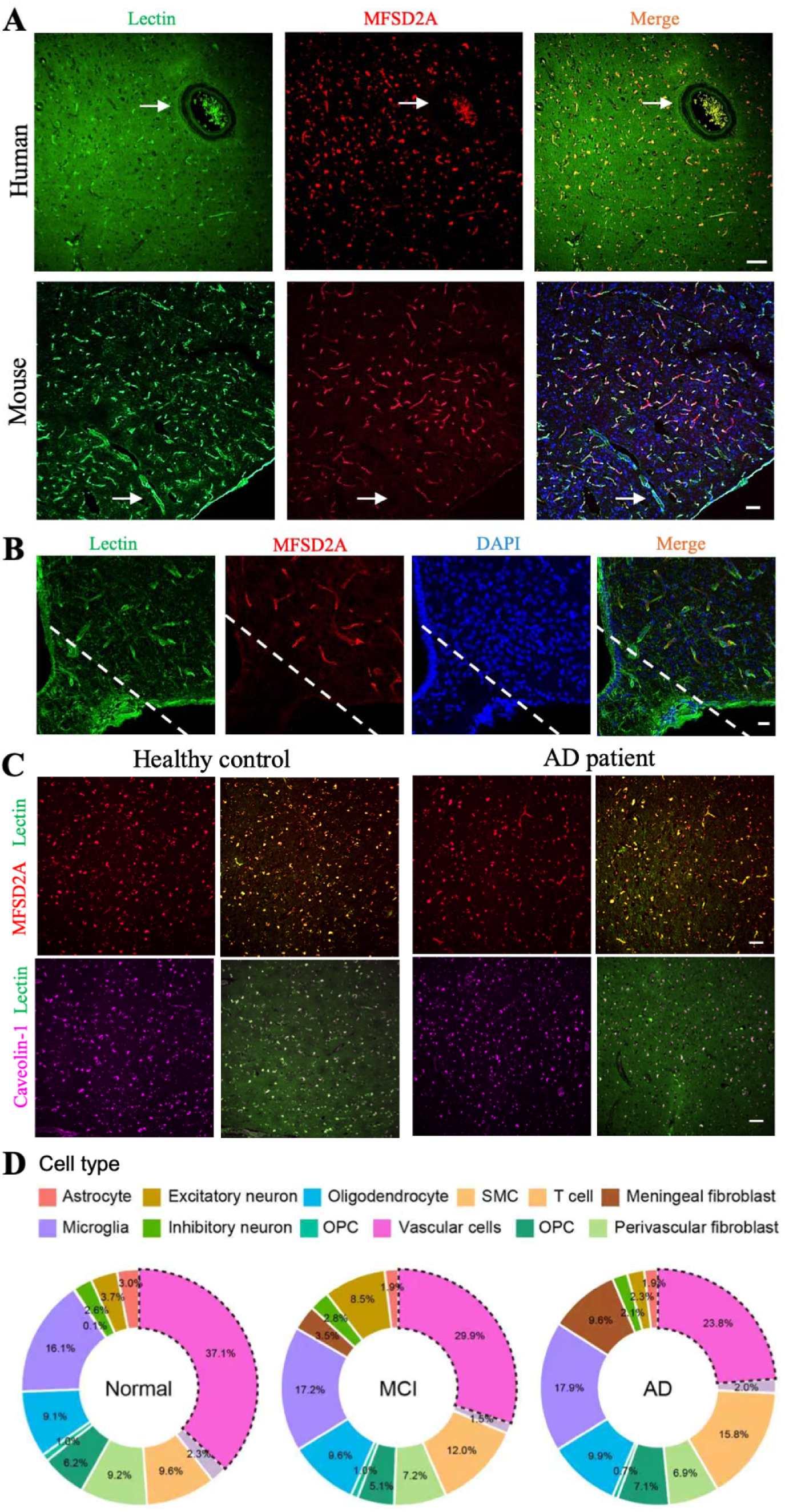
(A) Representative immunofluorescence images of MFSD2A and lectin-labelled vessels in the human and mouse brain. Arrows indicate large-calibre vessels with undetectable MFSD2A expression. (B) Representative immunofluorescence images showing undetectable MFSD2A expression in the arcuate nucleus and median eminence. Dashed lines delineate the boundary of the arcuate nucleus and median eminence. (C) Representative immunofluorescence images of MFSD2A, caveolin-1, and lectin-labelled vessels in the cortex of control and AD brains. (D) Relative proportions of major cell types identified by single-nucleus RNA sequencing in normal, mild cognitive impairment (MCI), and AD brains. Dashed outlines indicate the vascular cell population. Scale bars, 50μm (A), 20 μm (B), and 50 μm (C).

**Supplementary Figure 2.**
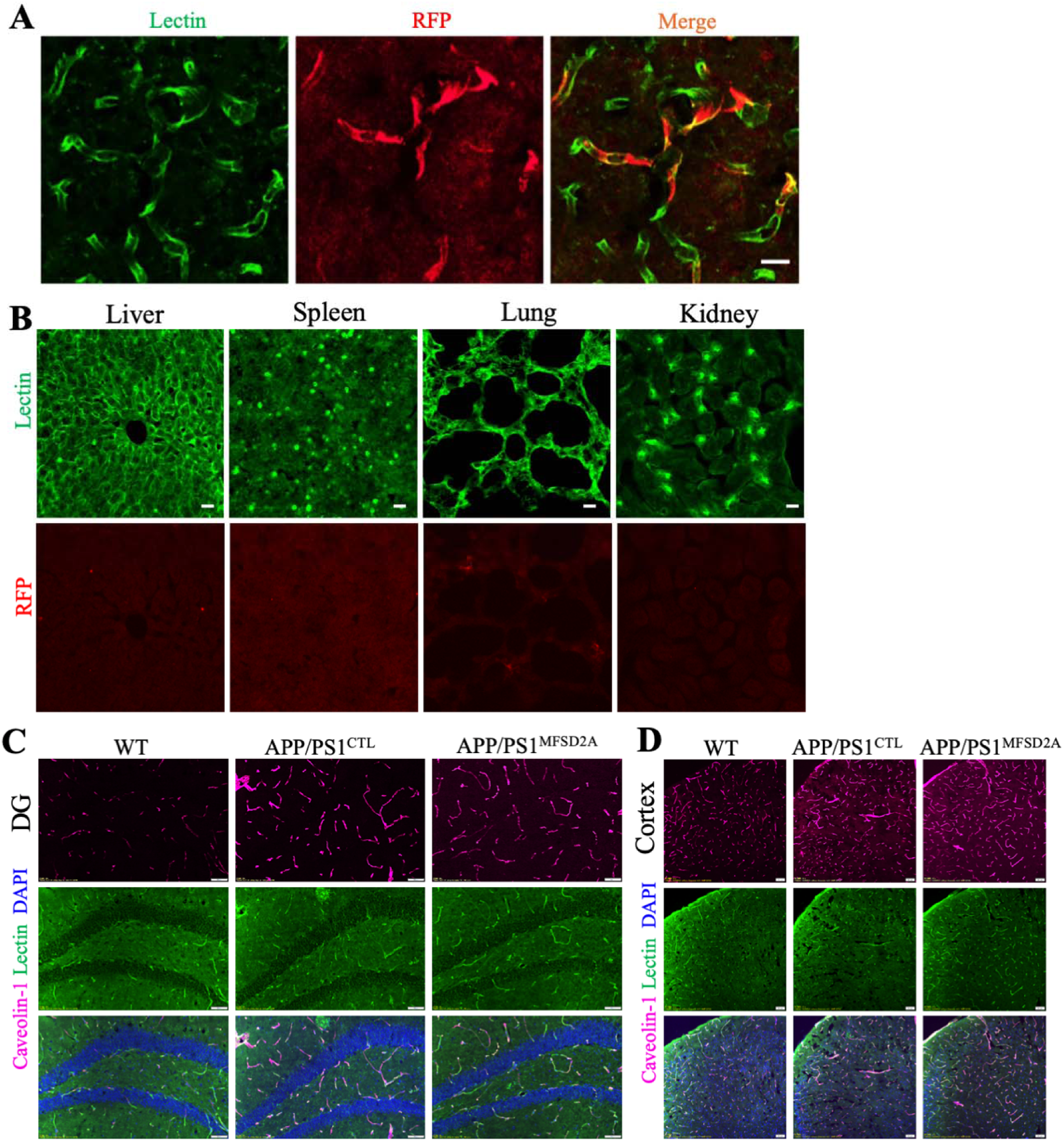
(A)Representative fluorescence images showing colocalisation of the AAV-BR1-encoded RFP reporter with lectin-labelled brain vessels. (B) Representative fluorescence images of lectin and RFP in the liver, spleen, lung, and kidney following systemic AAV-BR1 administration. (C and D) Representative immunofluorescence images of caveolin-1 and lectin-labelled vessels in the DG (C) and cortex (D) of WT, APP/PS1^CTL^, and APP/PS1^MFSD2A^ mice. Scale bars, 20 μm (A), 20 μm (B), and 50 μm (C and D).

**Supplementary Figure 3.**
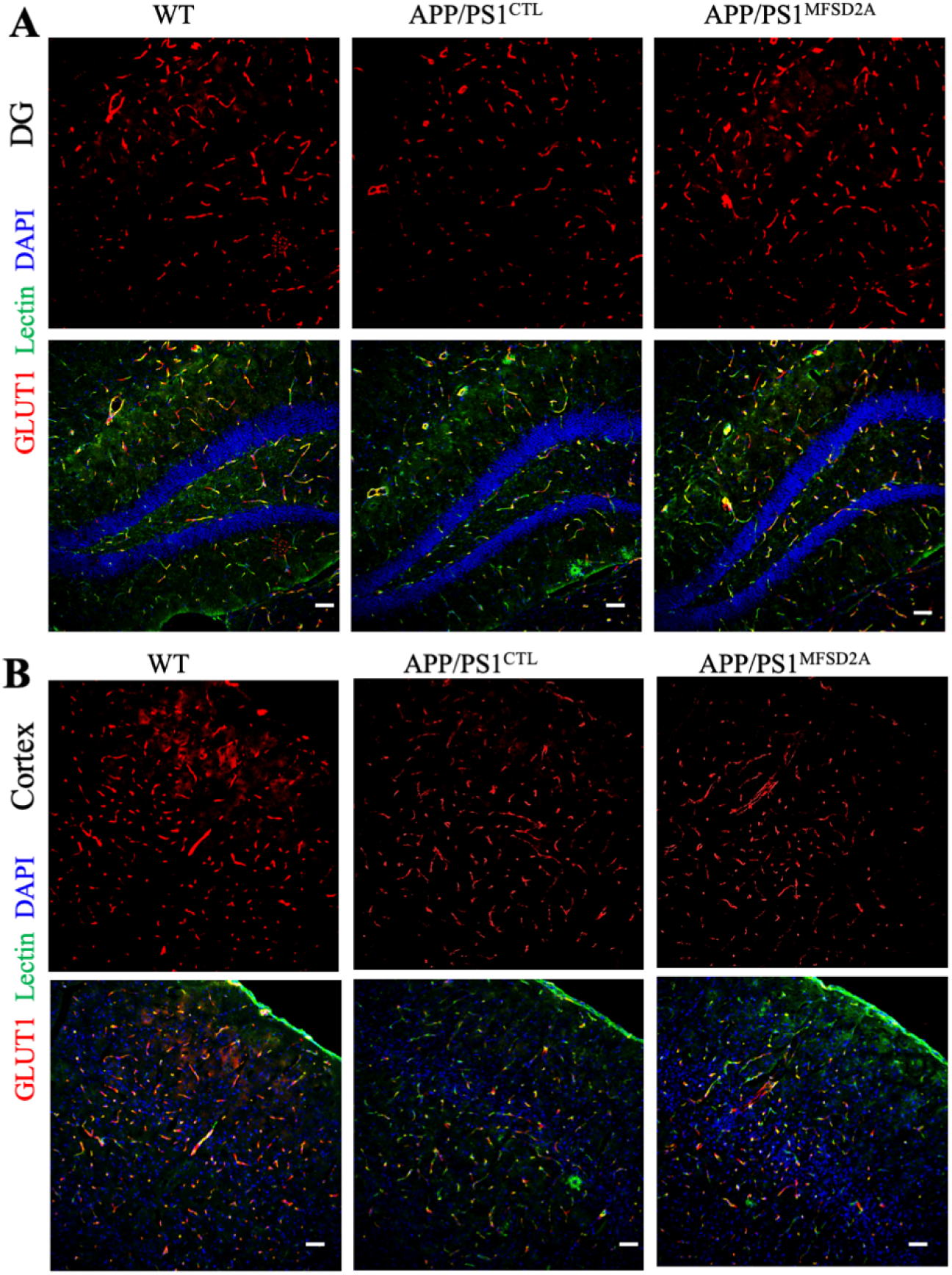
(A and B) Representative immunofluorescence images of GLUT1 and lectin-labelled vessels in the DG (A) and cortex (B) of WT, APP/PS1^CTL^, and APP/PS1^MFSD2A^ mice. Scale bars, 50 μm.

**Supplementary Figure 4.**
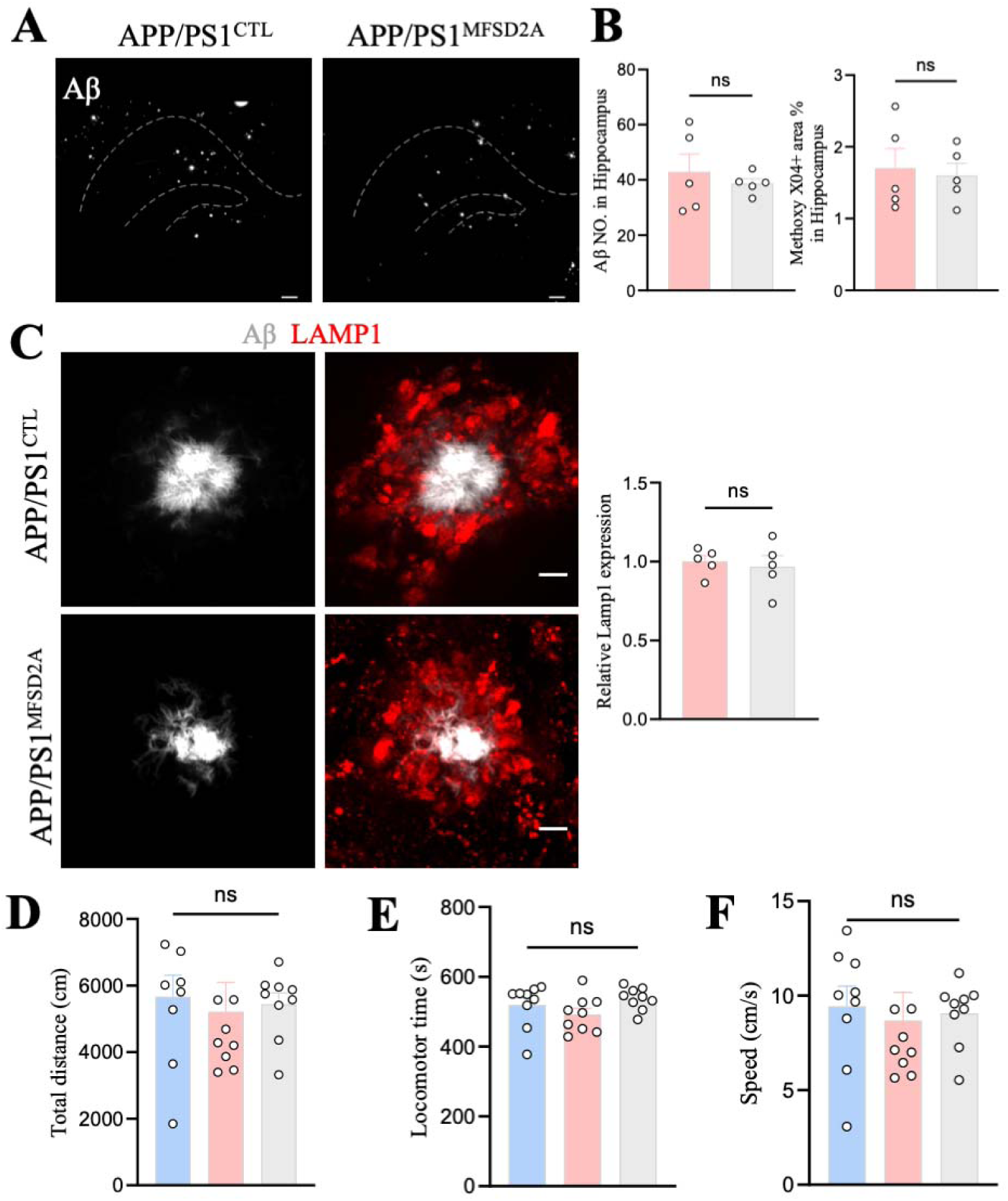
(A) Representative images of Methoxy-X04-labelled Aβ plaques in the hippocampus of APP/PS1^CTL^ and APP/PS1^MFSD2A^ mice. Dashed lines delineate the hippocampal region. (B) Quantification of hippocampal Aβ plaque number and plaque burden. (C) Representative immunofluorescence images and quantification of LAMP1 immunoreactivity surrounding Aβ plaques. (D-F) Quantification of locomotor time, total distance travelled, and average speed in the open-field test. Scale bars, 200 μm (A) and 5 μm (C). Data are presented as mean ± SEM; each dot represents one mouse. Statistical significance was assessed using an unpaired two-tailed Student’s *t*-test (B-C) or one-way ANOVA (D-F). ns, not significant.

**Supplementary Figure 5.**
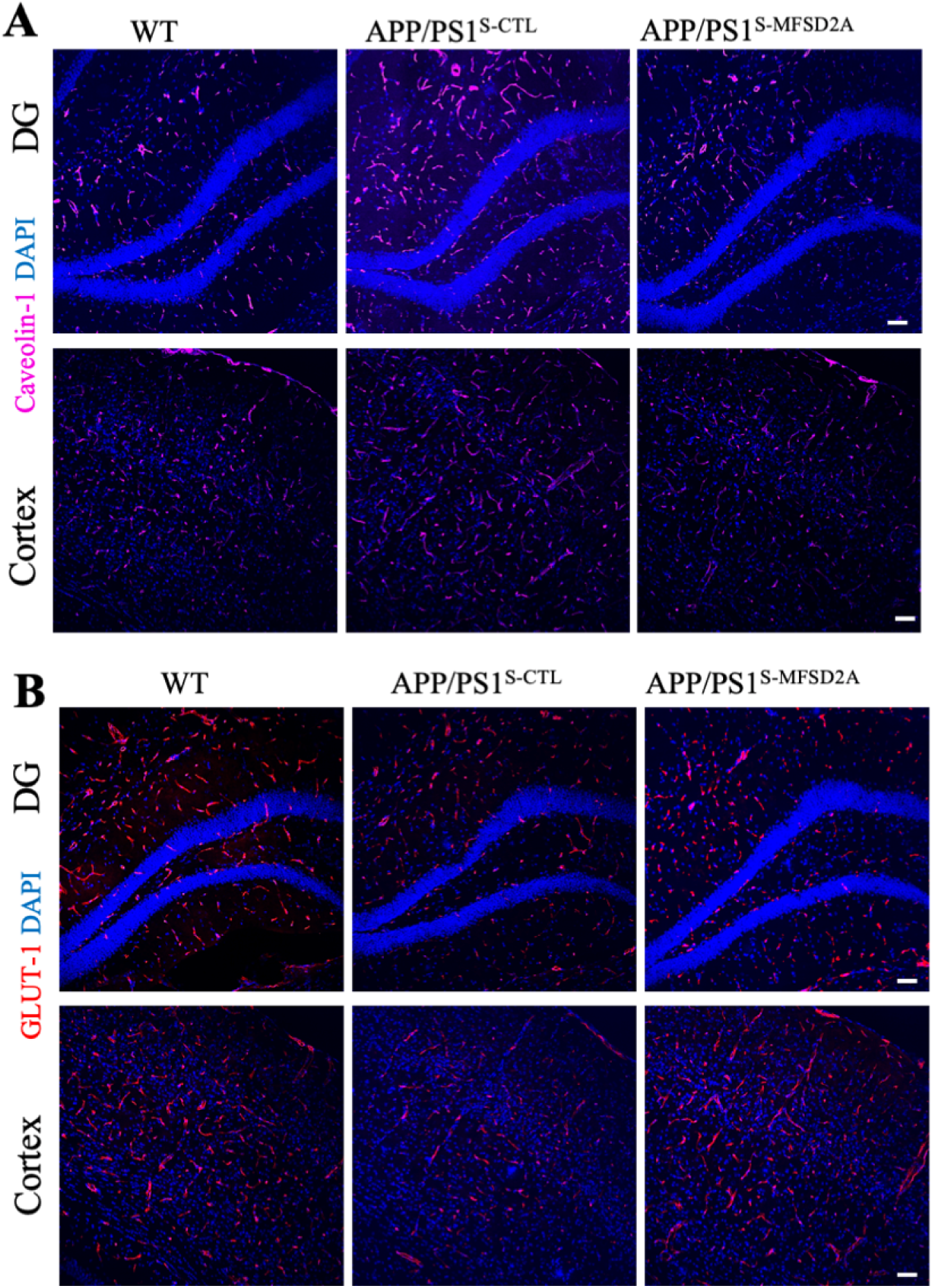
(A) Representative immunofluorescence images of caveolin-1 in the DG and cortex of WT, APP/PS1^S-CTL^, and APP/PS1^S-MFSD2A^ mice. (B) Representative immunofluorescence images of GLUT1 in the DG and cortex. Scale bars, 50 μm.

**Supplementary Table 1.** Information of healthy controls and AD patients.

| Information of healthy controls and AD patients |  |  |  |  |  |
| --- | --- | --- | --- | --- | --- |
| Group | Case | Age at death | Gender | Post-mortem delay (h) | ABC score for Level of AD |
| Control | 804 | 87 | F | 3.75 | Not |
| Control | 807 | 87 | M | 3.5 | Low |
| Control | 822 | 93 | M | 4.33 | Not |
| Control | 824 | 86 | F | 10.67 | Not |
| Control | 849 | 91 | F | 3 | Not |
| AD | 670 | 94 | M | 4.5 | High |
| AD | 709 | 92 | M | 5 | Intermediate |
| AD | 732 | 96 | M | 5.03 | Intermediate |
| AD | 768 | 94 | F | 4.8 | Intermediate |
| AD | 884 | 93 | M | 3.83 | High |

